# Prometheus: omics portals for interkingdom comparative genomic analyses

**DOI:** 10.1101/232298

**Authors:** Gunhwan Ko, Insu Jang, Namjin Koo, Seong-Jin Park, Sangho Oh, Min-Seo Kim, Jin-Hyuk Choi, Hyeongmin Kim, Young Mi Sim, Iksu Byeon, Pan-Gyu Kim, Kye Young Kim, Gukhee Han, Jong-Cheol Yoon, Yunji Hong, Kyung-Lok Mun, Banghyuk Lee, Deayeon Ko, Wangho Song, Yong-Min Kim

## Abstract

Functional analyses of genes are crucial for unveiling biological responses, for genetic engineering, and for developing new medicines. However, functional analyses have largely been restricted to model organisms, representing a major hurdle for functional studies and industrial applications. To resolve this, comparative genome analyses can be used to provide clues to gene functions as well as their evolutionary history. To this end, we present Prometheus (http://prometheus.kobic.re.kr),web-based omics portal that contains more than 17,215 sequences from prokaryotic and eukaryotic genomes. This portal supports interkingdom comparative analyses via a domain architecture-based gene identification system, Gene Search, and users can easily and rapidly identify single or entire gene sets in specific pathways. Bioinformatics tools for further analyses are provided in Prometheus or through BioExpress, a cloud-based bioinformatics analysis platform. Prometheus suggests a new paradigm for comparative analyses with large amounts of genomic information.

## Introduction

The completion of the Human Genome Project (2003) was not an end but rather a new beginning for further functional genomic analyses. The ENCyclopedia Of DNA Elements (ENCODE) was launched to begin investigating the functions of the identified human genes^1^. In addition, large-scale functional studies, such as interactome or network analyses, were performed in model organisms, including Arabidopsis thaliana, Saccharomyces cerevisiae, and Drosophila melanogaster. These efforts accumulated network information on various interactomes and gene functions. These vast amounts of biological information enabled functional studies that contributed to the unveiling of biological responses, the cloning of genes of interest, and the development of molecular markers for model organisms or medicines in humans^2,3^. Thus, the trend of functional analyses has been transferred from candidate gene research to genome-wide research. However, this flood of information has largely been restricted to model organisms, and it has been challenging for researchers to apply these data to newly sequenced genomes.

Since next-generation sequencing (NGS) technology was developed in the mid-2000s, an enormous amount of genomic information has been analyzed and amassed in public databases. As the numbers of sequenced genomes increased, many tools and pipelines were developed to investigate gene functions, identify gene families, and perform comparative genomic analyses. However, the application of comparative analyses is restricted to functional gene annotations and newly sequenced genome analyses. Newly sequenced genomes are initially compared to those that have previously been analyzed, including genomes of closely related species, to provide information on genome structure changes and gene repertoires. Such comparisons can also predict gene paralogues, which are genes related by duplication events, or orthologues, which are those related by speciation events^4-6^. As orthologues tend to be more similar in function that paralogues^7^, they are widely used for functional gene annotations^8^. Moreover, recent gene-of-interest studies that include multigenome orthologues offer insight into their mechanisms for adapting to the environment^9,10^. However, these comparative genomic analyses were performed at genome-, genus-, or kingdom-wide levels, thereby restricting comparisons to the species, family, or order level^11-13^. To understand the evolution of genes of interest more precisely, interkingdom analyses are needed, particularly because many genes in eukaryotic genomes have universal common ancestries in Bacteria and Archaea^14^.

Here, we report an omics portal for interkingdom comparative genomic analyses named Prometheus (http://prometheus.kobic.re.kr). We collected 17,215 sequences from 16,730 species and constructed four primary databases to provide basic genome information, with more detailed information on individual genes provided in secondary databases. Researchers can then access detailed information on genes of interest, such as gene structure, domain architecture, subcellular localization, orthologues, and paralogues, as well as their sequences. In particular, Prometheus provides Gene Search to identify genes of interest based on their domain architectures from prokaryotes to eukaryotes and performs various comparative analyses, such as comparison of chromosome sequences, sequence alignment, and phylogenetic analyses. Furthermore, researchers can perform various bioinformatics analyses with these and their own sequencing data in a cloud-based platform, BioExpress. Prometheus suggests a new paradigm for genome research, from single genes of interest to entire gene pathways.

## Materials and Methods

### Web interface

Prometheus furnishes data search, configuration of data analyses, data visualization, and storage of users’ own data. The interface is implemented using a Hypertext Markup Language (HTML), cascading style sheets (CSS) and uses a jQuery JavaScript library (jQuery) to modify web page contents. To visualize data, dynamic web interface is constructed by Asynchronous JavaScript and XML (Ajax) using JavaScript Object Notation (JSON) data format. Furthermore, genome browser was constructed using Scalable Vector Graphics (SVG) and phylogenetic viewer is constructed using JavaScript. Web interface of Prometheus supports a cross-browsing.

### Construction of taxonomy combined heatmap of photolyase/cryptochrome family

Sequences for the photolyase/cryptochrome family of genes from different species in previous study^15^ were collected and domain architectures were investigated using InterProScan v5.0^16^. Each of the subtypes reported in previous studies were investigated using Gene Search in Prometheus. The numbers of each of the subfamily genes were calculated for individual species and visualized as a heatmap using R scripts. The taxonomic tree was constructed using phyloT in iTOL^17^, an online tool that generates phylogenetic trees based on the NCBI taxonomy. Finally, the taxonomic tree and heatmap were combined using Adobe Illustrator.

### Bioinformatics analysis using a cloud-based analysis system, BioExpress

LAST^18^, BLAST^19^, Clustal Omega^20^, MUSCLE^21^ and InterPro^16^ programs are run in hybrid-cluster system, BioExpress. To support further genomic analyses using personal data such as RNA-Seq, Chip-Seq, or genome resequencing data, Prometheus links to BioExpress and users can perform further various genomic analyses using personal data in My Gene and various analysis pipelines in BioExpress. BioExpress is constructed by Hadoop to support high-speed analysis of a large amount of data. To maintain a large data of user, Prometheus stores the data divided by optimized block size using Hadoop Distributed File System (HDFS) into various computer servers. These storage system can maintain three copies of user data and provides stable data storage by reducing risk of data loss. Web server of Prometheus transmits task, progress and result of data analysis to BioExpress server using apache thrift library-based Remote Procedure Call (RPC) and received result as JSON format data. The result of genomic analyses is stored in HDFS and downloaded in web browser using HTTP protocol. In case of large amount of data, users can download their data using KoDS (KOBIC Data Transfer Solution). KoDS is a high-speed file transfer software using TCP/IP protocol and transferred user data is stored in HDFS.

### Construction of Database

The database consisted of primary and secondary data tables in the Prometheus was constructed using MySQL database management system. In database, primary data tables were created through data is opened in five public databases and secondary data tables were constructed by parsing results of bioinformatics tools such as InterProScan^16^, OrthoMCL^22^, MultiLoc2^23^ and TargetP^24^. Detailed methods for construction of database are described in Supplemental Note Section 1.

## Results

### Concept and construction of Prometheus

Prometheus provides an integrated pipeline for interkingdom comparative genomic analyses and comprises four major sections, Genome Archive, Gene Search, BioExpress, and Genome Analysis. Users can identify genes of interest using Gene Search and investigate their domain architectures using InterPro^16^ in Genome Analysis. Furthermore, users can obtain additional species information via accessing the Korean Bioresource Information System (KOBIS) or perform further analyses by accessing the cloud-based BioExpress (Figure 1).

**Figure 1.**
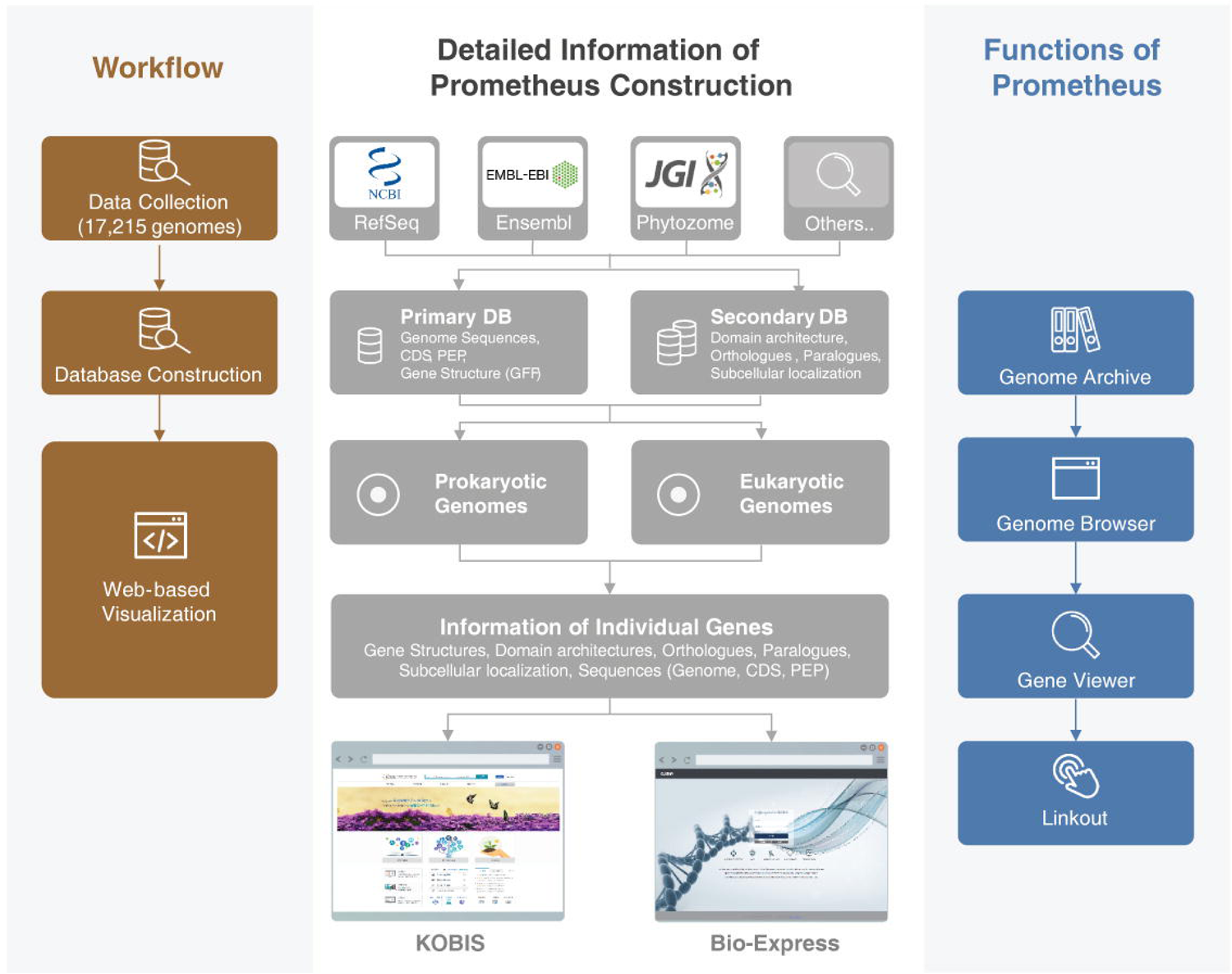
Concept and construction progresses of Prometheus. Schematic showing the workflow for the construction of Prometheus (left), detailed information for each stage (middle), and the functions available with Prometheus (right).

To establish Prometheus, 17,215 sequences from 16,730 species were collected and stored in four primary databases. The genomic information in Genome Archive (Figure 2A and Supplementary Table 1) is arranged by taxonomic rank (obtained from NCBI), which users can access by clicking the species name in the taxonomic tree or using a key word search. This General Information provides details on genome assembly, annotation, and taxonomy. In eukaryotic genomes, distinct versions of genome assembly and annotation were provided, and so each version is stored separately (Figure 2B). In prokaryotic genomes, genomic information is separated by strain to support metagenomics analyses. Genomes were classified according to criteria from RefSeq, which provided most of the genomic data (Supplementary Table 1), to construct the database and to visualize the genomic information. In total, 435 eukaryotic genomes, 15,984 prokaryotic genomes, and 311 archaea genomes were collected and assembled into the four primary databases containing information on assembled genomes, general feature formats (GFFs), coding sequences (CDSs), and protein sequences, for a grand total 213,478,449 records (Supplementary Table 2). Taxonomic information in Genome Archive is stored in a taxonomy database and general information of genome assembly and annotation is stored in a genome report databank. Totally, 51 database were constructed with 1,163,053,603 records (Supplementary Tables 3, 4, and 5).

**Figure 2.**
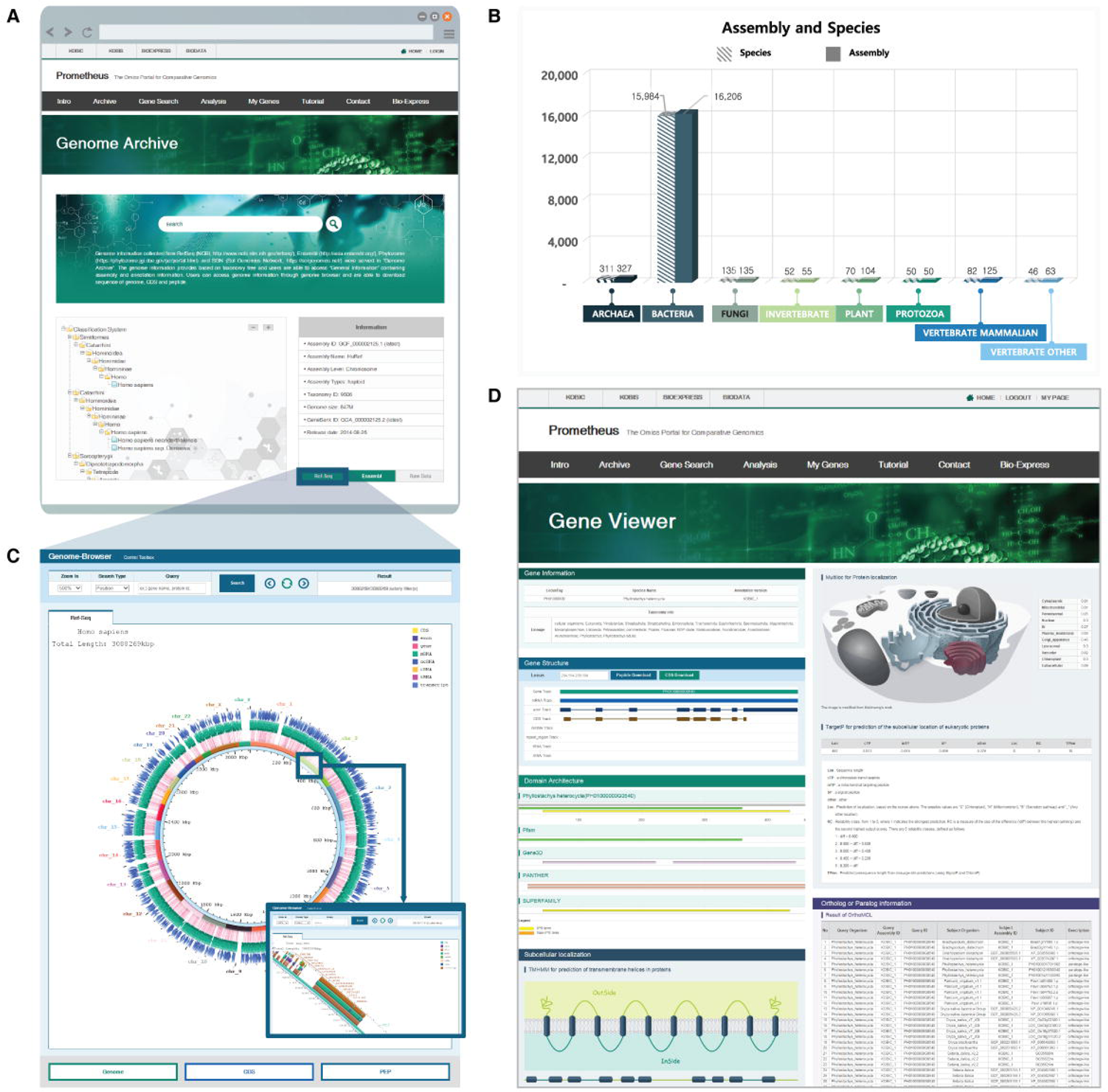
Construction of primary and secondary databases. (A) Screenshot of the Genome Archive page. (B) The numbers of species and genome versions used for the construction of Prometheus. (C) Screenshot of the Genome Browser of Prometheus. A region of the human genome (HGP 38) is shown enlarged in the small box. (D) Screenshot of the Gene Viewer for providing detailed information of individual genes. Gene structure, domain architecture, subcellular localization, and orthologous and paralogous genes are shown in each panel.

General information on individual genomes is obtained using Genome Browser (Figure 2C), with zoom in/out functions ranging from 100% to 1,000% and a gene search function by position or gene name. Users can access and download the individual gene’s information (CDS and peptide sequence) by key word search or by clicking within the Genome Browser. Detailed information on individual genes is provided in Gene Viewer (Figure 2D), and users can access the Genome Browser or result pages in Gene Search. Bioinformatics analyses, including InterPro^16^, OrthoMCL^22^, MultiLoc2^23^, and TargetP^24^ were performed using protein sequences of each species to construct six secondary databases, which are presented in separate sections within the Gene Viewer (Figure 2D). In summary, genomic information collected from major public databases is contained in the Genome Archive, and integrative information of genomes or individual genes from individual databases is accessed via Gene Viewer.

### Analyses of transcriptional factors and tricarboxylic acid (TCA) cycle in Gene Search

The major function of Prometheus is to perform interkingdom comparative analyses. To support this objective, secondary databases containing information on domain architectures and orthologues/paralogues of individual genes were constructed. We validated the utility of Prometheus by performing an interkingdom investigation of transcription factors (TFs) and genes involved in the TCA cycle using Gene Search (Figure 3, Supplementary Tables 6, and 7). A pipeline (iTAK v1.7)^25^ was used to identify plant TFs and classify protein kinases. TFs, transcriptional regulators (TRs), and kinases were identified by consensus rules mainly summarized from PlnTFDB^26^, PlantTFDB^18^ with families from PlantTFact^27^, and AtFDB^28^. Domain architectures of each TF were investigated using InterProScan, and their domain architectures depicted by InterPro entry (IPR) terms were used for further analyses using Gene Search. To provide additional information about identified genes, the number of domain subtypes are depicted in a summary table in Gene Search and as a header of sequence data in a FASTA file (Supplementary Figure 1). Users can categorize identified genes into each subtype. We identified and validated 79,960 genes from 15 gene families using the iTAK v1.7 (Figure 3A and Supplementary Table 6)^25^. The accuracy of our Gene Search ranged from 86.03% to 99.98%, with an average accuracy of 96.41%. High rates of accuracy were observed for genes encoding TFs containing significant IPR terms, such as FAR1, MADS-box, NAC or Dof domains, whereas those for TFs without significant IPR terms, such as B3-type TFs or CAMTA, showed lower rates. Thus, these data suggest that specific IPR terms or exact domain architectures are required to enhance the accuracy of Gene Search.

**Figure 3.**
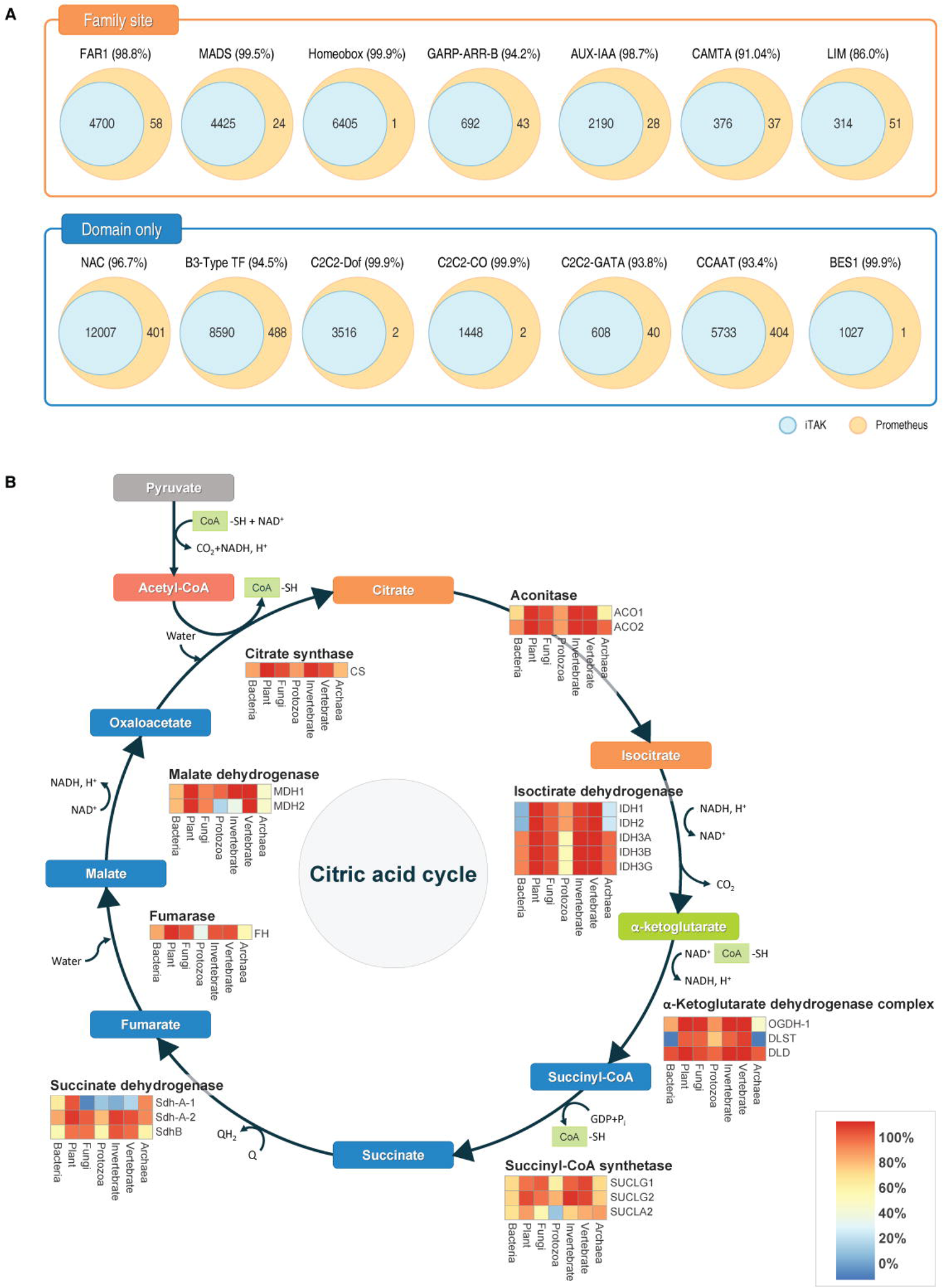
Identification of TFs and genes in the TCA using Gene Search. (A) Validation of identified TFs using iTAK pipeline. (B) Human TCA cycle genes were investigated and used for further analysis. The ratios of each gene are shown as heatmaps.

Genes involved in the TCA cycle were further investigated with Gene Search to demonstrate the potential for applying comparative genomics at the pathway level. As the TCA cycle is a fundamental metabolic pathway for survival in prokaryotes and eukaryotes, we selected this for an interkingdom comparative genomic analysis. A total of 435,044 genes were identified from 20 individual genes in the TCA cycle using Gene Search, and the ratios of species harboring each gene in the TCA cycle were shown as heatmaps (Figure 3B and Supplementary Table 7). These results showed that some genes, such as those encoding isocitrate dehydrogenase (IDH1 and IDH2) and malate dehydrogenase (MDH1 and MDH2) evolved in a lineage-specific manner. Furthermore, the results show the lineage-specific rates of functionally redundant genes, such as those encoding succinate dehydrogenase and succinyl-CoA synthase. This investigation of the TCA cycle also provided information on the gene repertoires and the evolution of the TCA cycle in each kingdom. Thus, Prometheus provides information for evolutionary studies of single genes or those in specific pathways, including the distributions and rates of genes, as well as repertoires of gene orthologues in pathways. In addition, Prometheus provides the domain architectures of genes as well as their CDSs and/or peptide sequences.

### Tools for comparative analyses and personalized management system via My Genes in Prometheus

To support comparative analyses in Prometheus, essential tools such as LAST (a program for comparing sequences at the chromosome level)^29^, Basic Local Alignment Search Tool (BLAST)^19^, and InterPro are provided in Genome Analysis (Supplementary Figure 2). Users can monitor the progress of analysis in a personalized page, My Genes (Supplementary Figure 3), and download the result files from each program via a file menu. In the case of data from InterProScan, the result file is shown in a graphic format and results are downloaded in a tsv file format (Supplementary Figure 4). Thus, users can investigate domain architectures of genes of interest and perform interkingdom identification using Gene Search.

We performed a comparative analysis of genes in the photolyase/cryptochrome family using a gene set from a previous study^15^ as a control (Figure 4 and Supplementary Table 8). The domain architectures of photolyase/cryptochrome subfamilies are the same and family IPR terms are different (Figure 4A), enabling a more accurate identification of each subfamily. The results also indicated lineage-specific distributions of photolyase/cryptochrome gene families in each kingdom. Furthermore, the gene repertoires of each subgroup of these families are shown in a combined taxonomy heatmap (Figure 4B), demonstrating lineage-specific evolution and the expansion of subgroups at the species level. These data demonstrate that Gene Search and bioinformatics tools in Genome Analysis in Prometheus support interkingdom comparative analyses. In summary, Prometheus provides the bioinformatics tools essential for comparative analyses, and users can combine these tools with interkingdom comparative analyses in Gene Search to unveil gene function or the evolution of genes/gene families.

**Figure 4.**
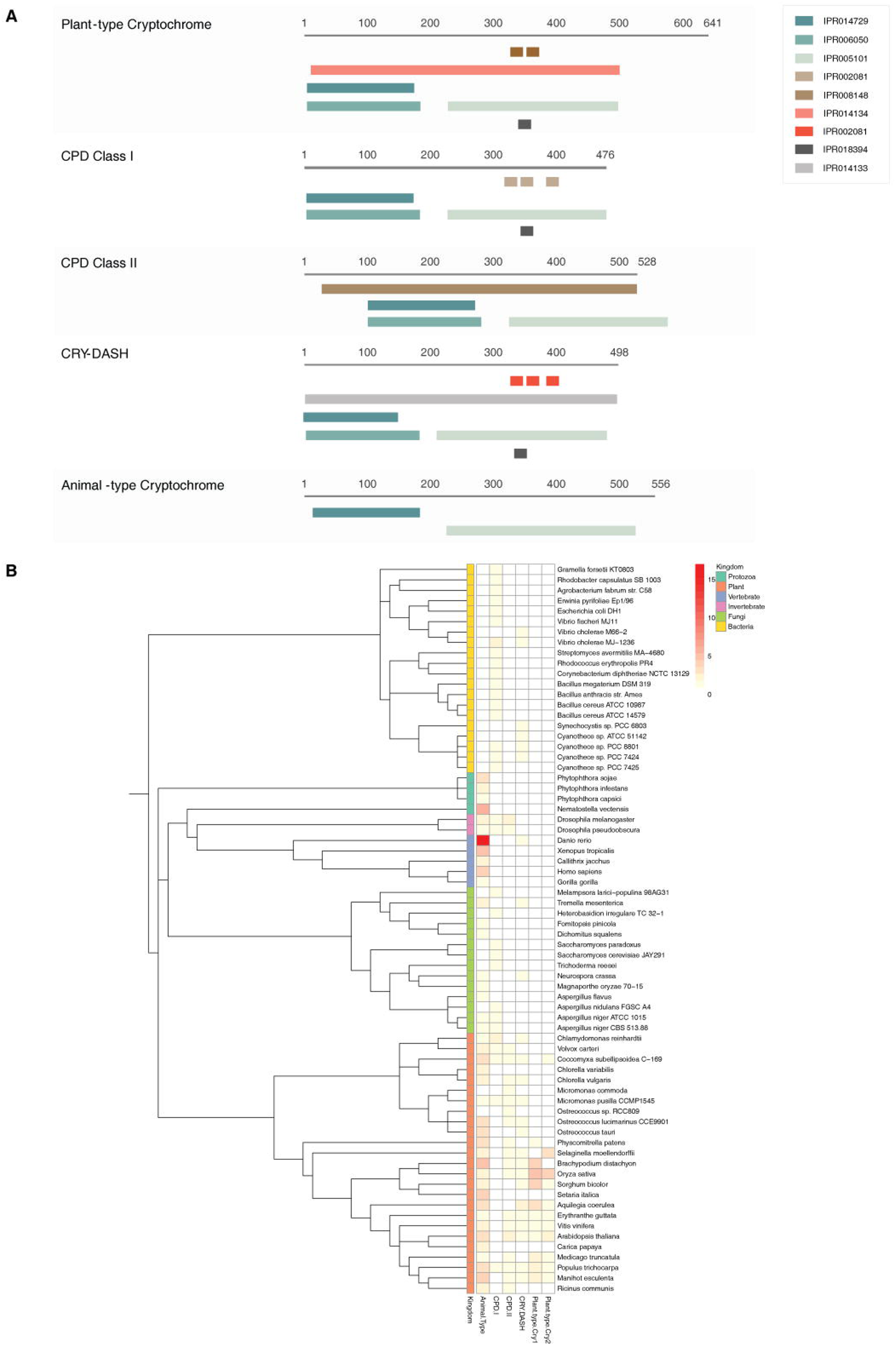
Interkingdom comparative analysis of the photolyase/cryptochrome gene family. (A) Domain architectures of the photolyase/cryptochrome gene family. (B) Distribution of photolyase/cryptochrome genes and taxonomic tree.

### Further genomic analyses using BioExpress with personalized data via My Genes

Personal data, such as RNA-Seq or Chip-Seq data, and downloaded data from Prometheus, such as genome sequences (genome, CDS, and peptide) in Genome Archive or FASTA files from Gene Search can be uploaded and stored in My Genes (Supplementary Figure 3) and further analyzed using the BioExpress platform (http://bioexpress.kobic.re.kr/bioexpress.en/). BioExpress is a cloud-based analysis platform, and programs for bioinformatics analysis are modularized and shown as icons (Supplementary Figure 5). Users can construct their own analysis pipelines by selecting and linking each modularized program using arrows.

We performed a transcriptomic analysis in BioExpress using the genome of *Hibiscus syriacus*^6^ and RNA-Seq data. For this, TopHat^30^ and Cufflink^31^ programs were used, and genes differentially expressed in tissues from a previous study were identified and visualized as a heatmap (Supplementary Figure 6). Thus, users can perform bioinformatics analyses with personal data in My Genes by linking to BioExpress. This combination of Prometheus and BioExpress can provide convenient and user-friendly analysis conditions for non-bioinformatician scientists.

## Discussion

Since NGS technology was developed and applied to biology, vast amounts of genomic data have accumulated. With these data, comparative analyses of species or genes can be performed to unveil gene function or evolution. For instance, the evolution of pungency in peppers was discovered by a comparative analysis with tomato and potato genomes^5^. However, only a small number of biologists can perform these comparative analyses using bioinformatics tools. Indeed, the accessibility of bioinformatics analysis is currently a major hurdle for ongoing biologic research. Thus, we constructed Prometheus, a web-based omics portal for interkingdom comparative genomic analyses. Biologists can identify genes or gene families of interest using the domain architectures in Gene Search. Genes from multigene families containing various domain architectures can be detected, such as for the photolyase/cryptochrome family^15^ and the nucleotide-binding leucine-rich repeat gene family^32^. Additional subtype information of identified genes is provided in the headers for their sequences in FASTA files.

The goal of combining kingdom-wide gene identification with subtype information is to provide evolutionary insight by detecting lineage-specific subtypes or subtype distribution patterns, as exemplified by the analysis of gene subtypes involved in the TCA cycle. Moreover, users can perform comparative analyses of single genes as well as sets of genes involved in specific signaling pathways. We found that genes containing specific domains showed high rates of accuracy in domain architecture-based Gene Search in Prometheus. However, the accuracy was reduced for genes without specific IPR terms, which is a limitation of domain architecture-based gene search systems using InterPro^16^ or the pfam^33^ database. Nevertheless, this limitation will be minimized as Prometheus is updated with new releases of these databases.

To support comparative analyses, Prometheus incorporates various tools, such as LAST^29^, Clustal Omega^20^, and Phylogeny viewer, in Genome Analysis. This is a valuable addition, as there are currently few web sites for comparative analyses with large restrictive or functionally important gene families, such as TFs. For TFs in plants, there are two representative web sites, PlnTFDB^26^ and PlantTFDB^18^, but their gene repertoires differ due to their rules for indemnification of TFs^25^. Prometheus clears this particular hurdle via its domain architecture-based Gene Search system, thereby providing biologists with a powerful comparative analysis platform with various tools for further studies.

Prometheus provides information to users on individual genomes assigned by taxonomy in Genome Archive via Genome Browser. Here, users can download the genomic and peptide sequences and CDSs as well as upload their own data for further analyses in Prometheus or the cloud-based BioExpress platform. Furthermore, user can access detailed information of genes of interest in the Gene Viewer page. The connection with BioExpress enables Prometheus to provide various bioinformatics tools and allows biologists to analyze their own data in same platform. Thus, unlike other comparative genomics portals or platforms, Prometheus provides tools not only for comparative analyses but also for genomic analyses, such as transcriptome or resequencing analyses. In conclusion, Prometheus is an integrated platform for interkingdom comparative genomic analyses with additional support for other genomic analyses with the user’s own data. Thus, Prometheus offers biologists a new paradigm for comparative genome analyses and evolution studies. The platform and InterPro version will be updated annually with newly sequenced genomes to ensure that broad and precise data are available to researchers. Furthermore, newly developed tools for comparative genomic analyses will continue to be added to support various analyses. Finally, visualization of domain subtype architectures identified by Gene Search is now being developed and will be available for updates in the near future.

## Availability of data and materials

The raw sequence reads of RNA-Seq from *Hibiscus syriacus* has been deposited at DDBJ/ENA/GenBank under accession SRP087036 (PRJNA341314). Detailed methods to perform comparative analysis of Prometheus are provided in Tutorial section (http://prometheus.kobic.re.kr)

## Acknowledgements

This work was supported by the Korean Research Institute of Bioscience and Biotechnology (KRIBB) initiative program and by a grant from the Ministry of Science, ICT, and Future Planning (MSIP) of the Korean government through the National Research Foundation (NRF-2010-0029345). We would like to express our gratitude to Seon-In. Yeom, Seungill Kim, Ah-Young Shin, and Rose Mikulski for their valuable comments during writing of this work.

